# Cysteine Reactivity Profiling Illuminates Monoclonal Antibody Disulfide Bond Reduction Mechanisms in Biopharmaceutical Process Intermediates

**DOI:** 10.1101/2025.04.16.649137

**Authors:** Taku Tsukidate, Zhenshu Wang, Andrew Hsieh, Patricia Rose, Xuanwen Li

## Abstract

Monoclonal antibodies (mAbs) are crucial biotherapeutics with increasing global demand, but their production can be impacted by the reduction of disulfide bonds. This study presents a chemical proteomics workflow aimed at elucidating the mechanisms underlying disulfide bond reduction in mAbs produced from Chinese hamster ovary (CHO) cells. We employed iodoacetamide-desthiobiotin (IA-DTB) and the parallel accumulation and serial fragmentation combined with data-independent acquisition (diaPASEF) methodology for cysteine reactivity profiling and successfully quantified approximately 4,500 cysteine-containing peptides from harvested cell culture fluids (HCCF). Our findings reveal that various protein disulfide oxidoreductases were active in reducing HCCF, offering critical insights into the redox environment affecting mAb stability. Notably, we quantified specific cysteine residues in enzymes such as glutaredoxin and thioredoxin domain–containing protein 12, suggesting potential links between their activity to disulfide bond dynamics. This workflow not only complements conventional abundance proteomics but also enhances the understanding of functional enzyme states in bioprocessing. Ultimately, our approach provides a promising strategy for enzymes contributing to disulfide bond reduction, paving the way for improved manufacturing processes of mAbs.

## INTRODUCTION

Monoclonal antibodies (mAbs) represent a pivotal class of biotherapeutics used in the treatment of various diseases, including cancer, multiple sclerosis, rheumatoid arthritis, lupus, and respiratory disorders. Since the U.S. Food and Drug Administration (FDA) approved the first mAb product in 1986, the development of commercial mAbs has rapidly accelerated. Notably, the global sales revenue for all mAb drugs alone totaled nearly $115.2 billion in 2018 and is expected to reach $300 billion by 2025, underscoring their integral role in contemporary medicine.^1,2^ Given the expanding role of antibody therapeutics in treating a diverse array of conditions, substantial emphasis has been placed on enhancing manufacturing processes to optimize both yield and product quality.^3,4^

Therapeutic mAbs are predominantly produced in mammalian cell systems, notably Chinese hamster ovary (CHO) cells, where the proper formation of disulfide bonds is crucial for the functional integrity of the antibody. In Immunoglobulin G (IgG) antibodies, two heavy and two light chains are connected by interchain disulfide bonds, which are more vulnerable to environmental factors compared to the intrachain bonds protected within the beta sandwich structure.^5,6^ The reduction of interchain disulfide bonds during the storage of harvest cell culture fluid (HCCF), which often spans several days prior to purification via affinity chromatography, presents considerable challenges. This situation not only threatens product quality but also results in increased purification complexity and a notable decrease in the stability of the therapeutic product.^7,8^

In response to these manufacturing challenges, the biopharmaceutical industry has increasingly focused on identifying underlying cause for disulfide bond reduction.^8^ For example, Trexler-Schmidt and co-workers have observed a correlation between disulfide bond reduction in harvested cell culture fluid (HCCF) and cell lysis and demonstrated that intracellular components can reduce mAbs.^9^ At the same time, Kao and co-workers have determined that these intracellular components were macromolecules based on a dialysis experiment and demonstrated that the thioredoxin system can reduce mAbs when biochemically reconstituted.^10^ Another study has later confirmed that thioredoxin 1 indeed contributes to mAb reduction by a genetic knockdown study.^11^ In addition, several studies have suggested that glycolytic enzymes fuel the thioredoxin system by generating NADPH in HCCF.^10,12,13^ Similarly, other studies have identified the glutathione system as another potential cause of mAb reduction.^14,15^ However, it is unclear whether these and potentially other protein disulfide oxidoreductases are active or directly reduce mAbs in HCCF.

To address these limitations in the proteome scale, we developed a chemical proteomics workflow and profiled cysteine reactivity in reducing and non-reducing HCCFs. We systematically optimized the workflow to achieve deep cysteinome coverage and quantified ∼ 4500 cysteine-containing peptides. Our unbiased chemical proteomic analysis suggested that several protein disulfide oxidoreductases were in their active states in reducing samples and complemented conventional abundance proteomics analysis and preceding studies. Collectively, we demonstrate that our chemical proteomics workflow represents a powerful method to advance mechanistic understanding of disulfide bond reduction.

## MATERIALS AND METHODS

### Protein samples

mAb1 cell broth and HCCF were internally sourced. The following commercially available proteins were mixed and used as internal standards: alcohol dehydrogenase (Sigma), carbonic anhydrase II (Sigma), myoglobin (Sigma), phosphorylase-b (Sigma), and enolase (Sigma). HeLa protein digest was purchased from Thermo Fisher Scientific.

### Cysteine reactivity profiling sample preparation

mAb1 HCCF (200 μL) was incubated with 2 mM iodoacetamide–desthiobiotin in DMSO (IA-DTB, MedChemExpress, 10 μL) at 25 °C for 1 h. Then 10 g/L Sera-Mag Speedbead Carboxylate-Modified [E3 & E7] (Cytiva, 100 μL) and acetonitrile (700 μL) were added and incubated at 25 °C for 10 min with 800 rpm agitation. The beads were washed three times with acetonitrile (1000 μL) and incubated with 50 mM tris-HCl pH 8.0 (1000 μL) containing 20 μg of trypsin (Promega) and 5 μg of LysC (Wako) at 37 °C overnight. The protein digest was collected in a new microcentrifuge tube. The beads were mixed with 50 mM tris-HCl pH 8.0 (500 μL) and the supernatant was combined with the protein digest. The protein digest was incubated with Pierce streptavidin magnetic beads (Thermo Scientific, 50 μL) at 25 °C for 2 h. The beads were washed with 0.1 % SDS in TBS (1000 μL × 3), TBS (1000 μL × 3), and water (1000 μL × 1). The enriched peptides were released by incubating the beads three times with 0.1 % formic acid in 50 % acetonitrile (50 μL) for 1 min with 800 rpm agitation, concentrated to dryness on a vacuum centrifuge, reconstituted with 0.015 % *N*-dodecyl-β-*D*-maltoside (DDM) (50 μL), and loaded on Evotip Pure (Evosep) according to the manufacturer’s instructions.

### Abundance proteomics sample preparation

mAb1 cell broth or HCCF (10 μL) were spiked with the mixture of five protein standards (1 μL) and incubated with 10 g/L Sera-Mag Speedbead Carboxylate-Modified [E3 & E7] (10 μL), water (10 μL) and acetonitrile (70 μL) at room temperature for 10 min with 800 rpm agitation.^16^ The beads were washed once with acetonitrile (200 μL) and incubated with 50 mM tris-HCl pH 8.0 (20 μL) containing 1 μg of trypsin and 0.25 μg of LysC at 37 °C overnight. The protein digest was diluted 100-fold with 0.1 % formic acid in water and loaded (20 μL) on Evotip Pure.

### LC–MS analysis

Peptides were separated on the Evosep One LC system^17^ with Pepsep C18 columns (Bruker) using the manufacturer’s pre-defined methods denoted by daily sample throughput e.g., 30 samples per day (SPD) and analyzed on the Bruker timsTOF Pro 2 system. The standard DDA PASEF method i.e., “DDA PASEF-standard_1.1sec_cycletime.m” was used to build a spectral library. The diaPASEF method for cysteine reactivity profiling was optimized with the py_diAID software^18^ to target IA-DTB–modified precursors in the spectral library and covered 400–1300 m/z with 25 diaPASEF scans and two ion mobility windows per diaPASEF scan (i.e., 50 variable windows) (**Supporting Information Figure 1** and **Supporting Information Table 1**). The diaPASEF method for abundance proteomics was previously described^19^ (**Supporting Information Figure 2** and **Supporting Information Table 2**). In all methods, the ion mobility range was set as 0.7–1.43 Vs cm^−2^ with 100 ms of accumulation time and 100 ms of ramp time unless otherwise noted.

### Cysteine reactivity profiling data analysis

DDA PASEF data were processed with MaxQuant v2.6.11^20^. Peptide lengths of at least 7 amino acids, with maximum peptide mass of 4600 Da, and with up to two missed cleavages were permitted. The following variable modifications were applied: methionine oxidation, N-terminus acetylation, and the IA-DTB modification on cysteine (+296.1848406575 Da). The PSM FDR was set at 0.01. The evidence file and the msms file were filtered to remove reverse hits and joined and formatted as a single tab-separated table (.tsv) with a custom R script (**Supporting Information Script 1**). This .tsv file was fed to DIA-NN v1.9^21^ and reformatted, declaring additional options to specify library headers and to apply the IA-DTB variable modification. Subsequently, diaPASEF data were searched against this spectral library. The IA-DTB variable modification was applied. Mass accuracies were set at 15 ppm. Scan window was fixed at 9. Peptidoforms scoring and match-between runs were enabled. The QuantUMS algorithm^22^ was used in quantification. The parquet report was filtered requiring Lib.Q.Value < 0.01, Lib.PG.Q.Value < 0.01, Lib.Peptidoform.Q.Value < 0.01, Lib.PTM.Site.Confidence > 0.99, tryptic end, and the IA-DTB modification. Contaminant and decoy hits were removed. Peptide intensity was represented by the Precursor.Normalised value of the most intense precursor associated with the peptide. The sequence database contained the in-house CHO cell protein sequences, mAb1, spiked-in protein standards, and common contaminants.

### Abundance proteomics data analysis

diaPASEF data were processed with DIA-NN v1.9 in the library-free mode. A spectral library was built *in silico* for precursors with the mass-to-charge ratio between 300–1200 *m*/*z*, with minimum charge of two, and with up to one missed cleavage. N-terminal methionine excision was enabled. No other modification was applied. Subsequently, raw files were searched with this spectral library. Mass accuracies were set at 15 ppm. Scan window was fixed at 9. Match-between runs were enabled. The legacy (direct) method was used in quantification. Normalization was turned off. Precursor.Normalised values were log2-transformed and normalized based on the spiked-in protein standards. Precursors were filtered by requiring quantification in all sample pools, coefficient of variation across sample pools < 0.5, Lib.Q.Value < 0.01, and Lib.PG.Q.Value < 0.01. Protein quantity was calculated from the log2-transformed and normalized Precursor.Normalised values with the fast_MaxLFQ function implemented in the iq package^23^. The sequence database contained the in-house CHO cell protein sequences, mAb1, spiked-in protein standards, and common contaminants.

### HCCF Staging and Non-reducing capillary electrophoresis sodium dodecyl sulfate (NR CE-SDS)

HCCF was placed in an airtight bag to replicate the manufacturing scale of the HCCF holding. At each time point, a sample was taken and immediately spiked with N-ethylmaleimide after being withdrawn from the bag.^24^ Sample purity testing was performed using high-throughput analysis on the Perkin Elmer LabChip® GXII Touch Capillary Electrophoresis System as described by the vendor. Briefly, the preparation of the non-reducing buffer containing 1 M Iodoacetamide was prepared using 21 µL of 1 M Iodoacetamide with 700 µL non-reducing buffer (HT Protein Express Sample Buffer). The sample was prepared by mixing a small volume (2 µL) of protein with a 7 µL non-reducing sample buffer mixture. The protein mixture was centrifuged at 1500 g for 1 min and heated at 70 °C for 10 min. After heating, 35 µL of water was added to the mixture, centrifuged at 1,500 g for 5 min, and the samples were run under non-reducing conditions. The non-reduced protein was electrokinetically loaded directly into the chip from a microtiter plate (Bio-Rad Laboratories) using the separation channel from the chip.

## RESULTS AND DISCUSSION

### Functional Proteomics Approach towards Elucidating mAb Disulfide Reduction Mechanisms

Protein disulfide oxidoreductases are redox enzymes that catalyze dithiol-disulfide exchange reactions with a CXXC sequence motif at their active site.^25^ In the first step of a dithiol-disulfide exchange reaction between a disulfide oxidoreductase and another protein, the N-terminal active site cysteine attacks the disulfide bond of the substrate and forms a mixed disulfide. Next, the C-terminal active site cysteine attacks the mixed disulfide and releases the reduced substrate. Finally, the remaining disulfide between the active site cysteines is reduced with NADPH by thioredoxin reductase or glutathione reductase and glutathione to complete a cycle. Thus, an electrophilic chemical reporter targeting the active site cysteine can provide the basis for a quantitative readout of the functional state of individual disulfide oxidoreductases.^26,27^ To this end, we employed the cysteine-reactive chemical reporter iodoacetamide-desthiobiotin (IA-DTB)^28^ to selectively enrich peptides that contain highly nucleophilic cysteines from HCCF digests (**Figure 1**). HCCF samples were treated with 100 μM IA-DTB and proteins were aggregated and captured on beads^16^ to remove excess IA-DTB prior to on-bead digestion. Desthiobiotin-modified peptides were enriched on streptavidin beads and released for LC-IMS-MS/MS analysis. To achieve deep proteome coverage and precise quantification, we adopted parallel accumulation–serial fragmentation combined with data-independent acquisition (diaPASEF)^29^ in our workflow. Initially, a spectral library of IA-DTB-modified precursors were built with a longer (88 min, 15 SPD) chromatographic gradient and a data-dependent acquisition (DDA) method^30^. Subsequently, samples were analyzed using a shorter (44 min, 30 SPD) chromatographic gradient with a data-independent acquisition method, and raw data were searched against the spectral library.

**Figure 1.**
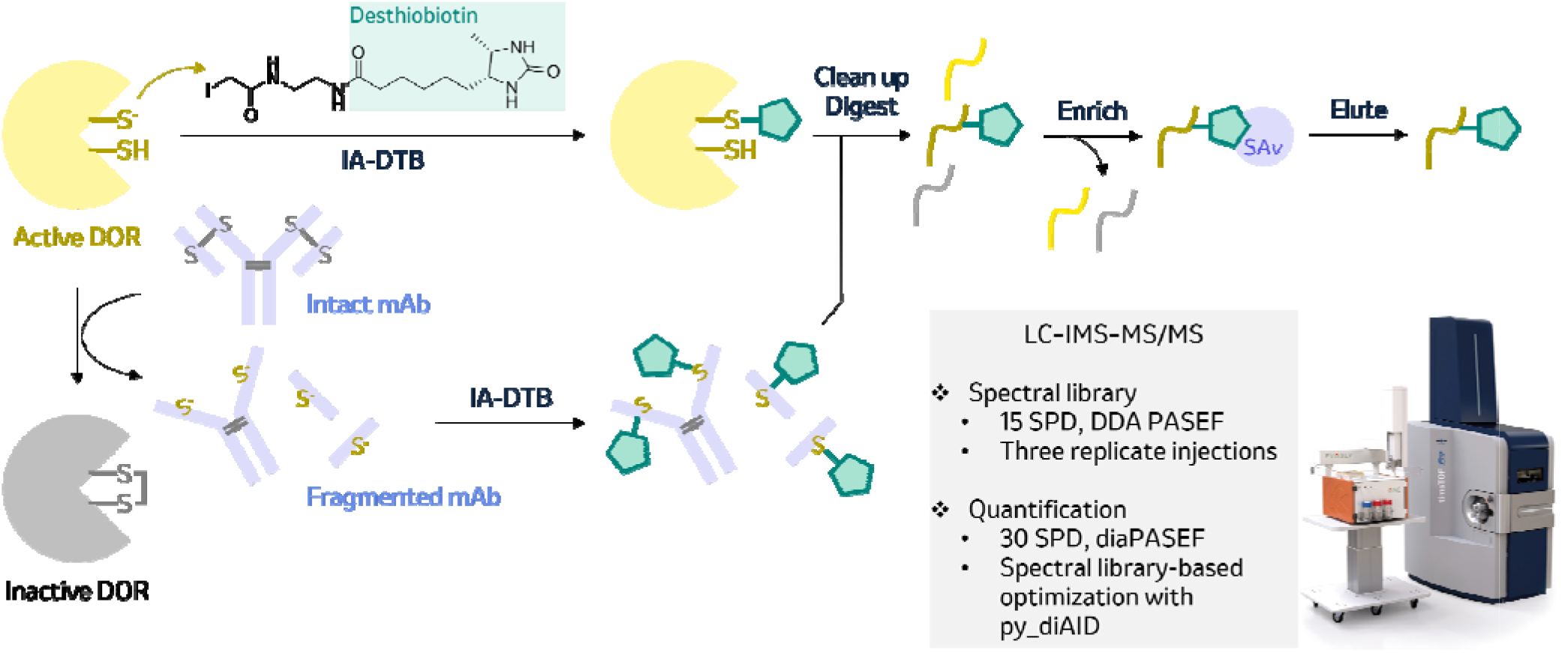
Schematic illustration of the cysteine reactivity profiling workflow to assess the functional state of individual disulfide oxidoreductases (DORs) in biopharmaceutical downstream process intermediates. IA-DTB labels reactive cysteine residues in the active site of DORs and on other proteins including mAb. Excess IA-DTB is removed prior to protein digestion. IA-DTB-modified peptides are enriched on streptavidin (SAv) beads and eluted for liquid chromatography–ion mobility spectrometry–tandem mass spectrometry (LC-IMS-MS/MS) analysis.

### Development of Cysteine Reactivity Profiling Workflow

To enable in-depth cysteine reactivity profiling, we systematically investigated and optimized key steps in the workflow. First, we evaluated the amount of starting protein input material (**Figure 2A**). Processing large amounts of material can be expensive and cumbersome even though material availability is typically not a limiting factor in host cell protein (HCP) analysis. Thus, we varied the amount of starting protein input material between 500 and 2000 μg and analyzed these samples with a longer gradient (88 min) and a DDA method. The number of IA-DTB-modified peptides modestly increased by ∼ 600 from 500 to 1000 μg and thereafter reached near saturation. Next, we evaluated offline sample fractionation to build a comprehensive spectral library (**Figure 2B**). On one hand, we processed 6000 μg protein input material as described above, subjected the IA-DTB-modified peptides to high-pH reversed-phase fractionation (HpHRPF) on a spin column to yield ten fractions, and analyzed each fraction with a longer gradient (88 min) and a DDA method. On the other hand, we processed and analyzed 2000 μg protein input material three times. The repeated injection method identified more IA-DTB-modified peptides than the HpHRPF method, perhaps by minimizing sample loss, and most identifications was shared between the two methods. Previous study has also demonstrated the advantage of building a spectral library without fractionation.^31^ Finally, we optimized the diaPASEF window design (**Figure 2C**). On average, IA-DTB-modified precursors had larger mass-to-charge ratios than the precursors that were identified in HeLa protein digest due to the addition of the desthiobiotin tag (+296.1848 Da). Thus, we generated a new window design on py_diAID^18^ focusing on 400–1300 *m*/*z* to target IA-DTB-modified precursors efficiently.

**Figure 2.**
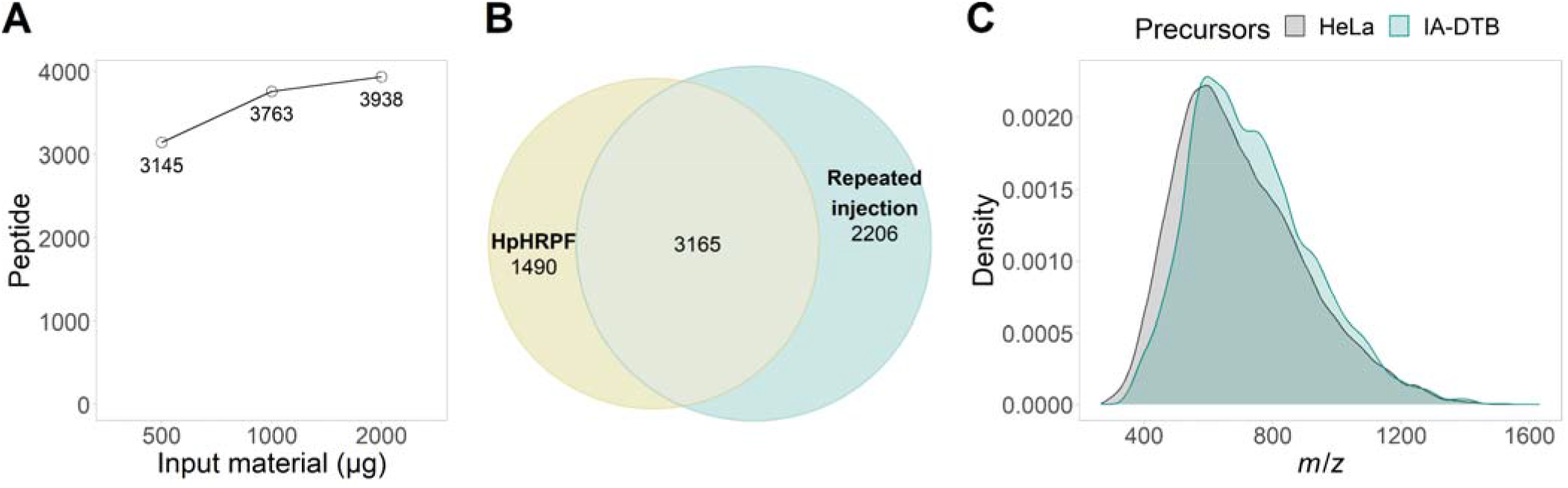
Development of the cysteine reactivity profiling workflow. (A) Evaluation of starting protein input material. (B) Evaluation of offline sample fractionation. (C) Comparison of precursor mass-to-charge ratio distribution. mAb1 HCCF. 15 SPD. DDA PASEF.

### Cysteine Reactivity Profiling Uncovers Reactive and Functional Cysteines in Disulfide Oxidoreductases

Next, we applied the cysteine reactivity profiling workflow to HCCF samples that were incubated for several days and up to two weeks. CHO cell culture for mAb1 was depth-filtered and samples were collected throughout the filtration. A late fraction that corresponds to high differential pressure i.e., strong resistance to flow across the filter was incubated at 4 °C and mAb1 reduction was monitored with the non-reducing capillary electrophoresis sodium dodecyl sulfate (NR CE-SDS) assay over two weeks (**Figure 3A**). The mAb1 main peak decreased and the low molecular weight species peak increased over the first three days, which was followed by a gradual reversing trend over next six days, indicating initial interchain disulfide bond reduction and subsequent re-oxidization. Thus, we profiled cysteine reactivity in samples taken at days zero, three, six and fourteen that correspond to different stages of mAb reduction and oxidation. Overall, we identified ∼ 4500 cysteine-containing peptides derived from ∼ 2300 proteins including mAb1 (**Figure 3B**). For example, we quantified the mAb1 heavy chain–derived peptide containing the cysteine residue that forms the interchain disulfide bond between light chain and heavy chain (**Figure 3C**). Its intensity increased from day zero to day three and decreased thereafter on days six and fourteen, reflecting the reduction and re-formation of this disulfide bond. In addition, we quantified 21 cysteine residues on 12 disulfide oxidoreductases including four active site residues with CXXC domain which might affect the mAb’s redox stability (**Figure 3D**). For example, we identified the N-terminal active site cysteine of glutaredoxin (GLRX, C23), which initiates a dithiol-disulfide exchange reaction by a nucleophilic attack on the substrate disulfide bond^25^, and observed a decrease in its intensity on day six and onwards (**Figure 3E**). The C-terminal active site cysteine of GLRX (C26) was identified only in the presence of the C23 modification, perhaps reflecting on its attenuated reactivity compared to C23. The allosteric regulatory cysteine residue C8 is sensitive to oxidation and contributes to oxidative inactivation of GLRX.^32^ This residue also maintained its reactivity on days zero and three, suggesting that the enzyme was not inactivated through this mechanism during first several days. Previous studies have implicated GLRX in mAb reduction but have been limited by the lack of selective assay, selective inhibitor, and active recombinant enzyme.^10,14,15^ Here, we demonstrated that GLRX were indeed in the active state in mAb-reducing HCCF. In another example, we identified the N-terminal active site cysteine of thioredoxin domain–containing protein 12 (TXNDC12, C64). TXNDC12 has the redox potential of ∼ −165 mV which is comparable to its native environment endoplasmic reticulum (−135 to −185 mV) and can function as both oxidase and reductase in cells.^24^ The decreasing active site thiol population suggests reductase activity in our samples. In contrast, quiescin sulfhydryl oxidase 1 (QSOX1) is an oxidase that contains a thioredoxin (Trx) domain and an essential for respiratory and vegetative growth (Erv) domain as well as a non-covalently bound flavin adenine dinucleotide (FAD) cofactor.^33^ These two domains relay electrons from a dithiol-containing protein substrate to an electron acceptor such as oxygen, introducing a disulfide bond to the protein substrate. The C456 residue corresponds to the C-terminal active site cysteine in the Erv domain. Its intensity peaked on day six, which coincided with mAb re-oxidation. Taken together, these results demonstrate that the cysteine reactivity profiling can provide a quantitative readout of the activity state of individual disulfide oxidoreductases.

**Figure 3.**
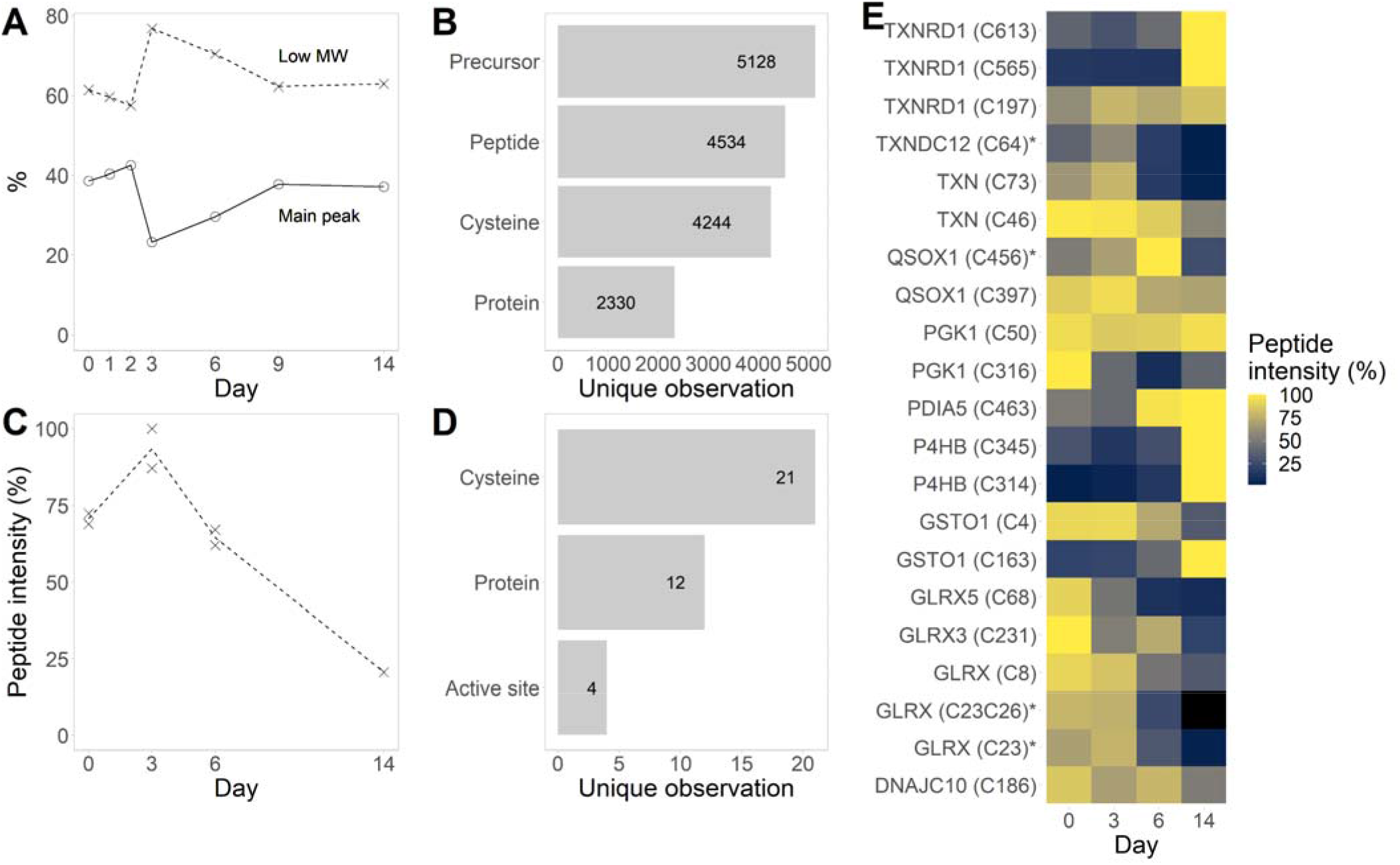
Cysteine reactivity profiling in disulfide-reducing and thiol-oxidizing HCCFs. (A) mAb purity monitored at room temperature over two weeks. NR CE-SDS. (B) Depth of cysteine reactivity profiling. (C) Intensity of the mAb1 heavy chain–derived peptide carrying an IA-DTB-modification on the cysteine residue that forms the interchain disulfide bond between light chain and heavy chain. (D) Coverage of disulfide oxidoreductases. (E) Average peptide intensities of disulfide oxidoreductase–derived peptides. Asterisks indicate active site cysteines. 30 SPD. diaPASEF. N = 2.

### Cysteine Reactivity Profiling Complement Conventional Abundance Proteomics

Finally, we assessed the cysteine reactivity profiling in comparison to conventional abundance proteomics. Thus, we clarified mAb1 cell broth with either centrifugation or depth filtration, processed these samples for abundance proteomics analysis with the protein aggregation capture method^16^, and analyzed them with the diaPASEF method as we previously described^19,29^. mAb1 was stable at room temperature in cell broth but not in depth-filtered HCCF, suggesting cell lysis and concomitant release of intracellular proteins and/or other metabolites during depth-filtration promotes mAb reduction^9,10,15^ (**Figure 4A**). We quantified 4470 HCPs including 26 disulfide oxidoreductases in the proteome analysis, which far exceeds the proteome coverage in similar studies and can provide more comprehensive view^12,13^ (**Figure 4B**). Consensus clustering analysis grouped these HCPs into four clusters. Most proteins belonged to the cluster one and were more abundant in depth-filtered HCCF, especially in fractions that were collected with high differential pressure, than those in cell broth or centrifuged HCCF. Subcellular localization analysis revealed that this cluster was enriched with cytoplasmic and nuclear proteins, confirming that many intracellular proteins were released during depth filtration presumably through cell lysis (**Figure 4C**). Indeed, TXN and TXN reductase 1 (TXNRD1) (cluster one)^10,11,14^ as well as glutathione reductase (GSR, cluster three)^14^, which have been implicated in mAb reduction, were more abundant in depth-filtered harvest. In contrast, HCPs in the cluster four appeared to be more abundant in cell broth and centrifuged harvest and this cluster was enriched with extracellular i.e., secreted proteins. For example, protein disulfide isomerase A4 (PDIA4) localizes in endoplasmic reticulum and/or is secreted and may catalyze disulfide bond formation in mAb.^34^ In addition, QSOX1 QSOX2 function as disulfide oxidases. These results demonstrate that abundance proteomics quantify thousands of HCPs in HCCF and connect mAb reduction to individual HCP abundance. and can

**Figure 4.**
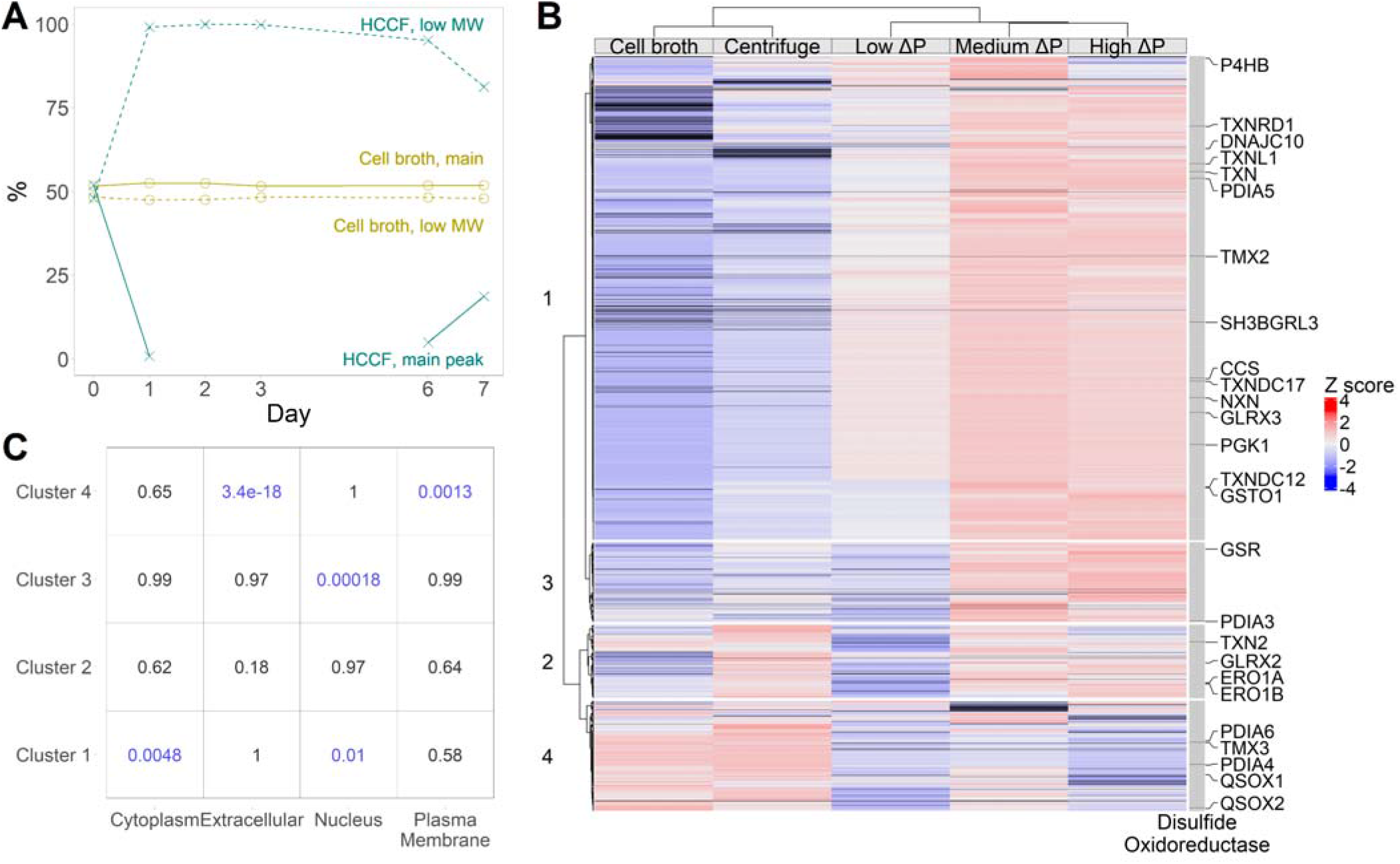
Abundance proteomics in stable cell broth and disulfide-reducing HCCFs. (A) mAb purity monitored at room temperature over seven days by NR CE-SDS. (B) Heatmap of relative protein abundance. ΔP: differential pressure. (C) Subcellular localization analysis of consensus clusters. Numbers indicate p values from Fisher’s exact test.

However, abundance proteomics cannot distinguish activity states of individual enzymes. In this regard, the cysteine reactivity profiling has unique advantage as a functional proteomics method and can provide complementary information (**Figure 5**). For example, the cysteine reactivity profiling revealed that GLRX and TXNDC12 were in the dithiol state which could reduce a disulfide bond–containing protein substrate in mAb-reducing samples. Moreover, the cysteine reactivity profiling uniquely identified GLRX, perhaps thanks to improved sensitivity afforded by peptide enrichment. Nevertheless, the cysteine reactivity profiling provided the evidence to demonstrate these enzymes were active in our samples.

**Figure 5.**
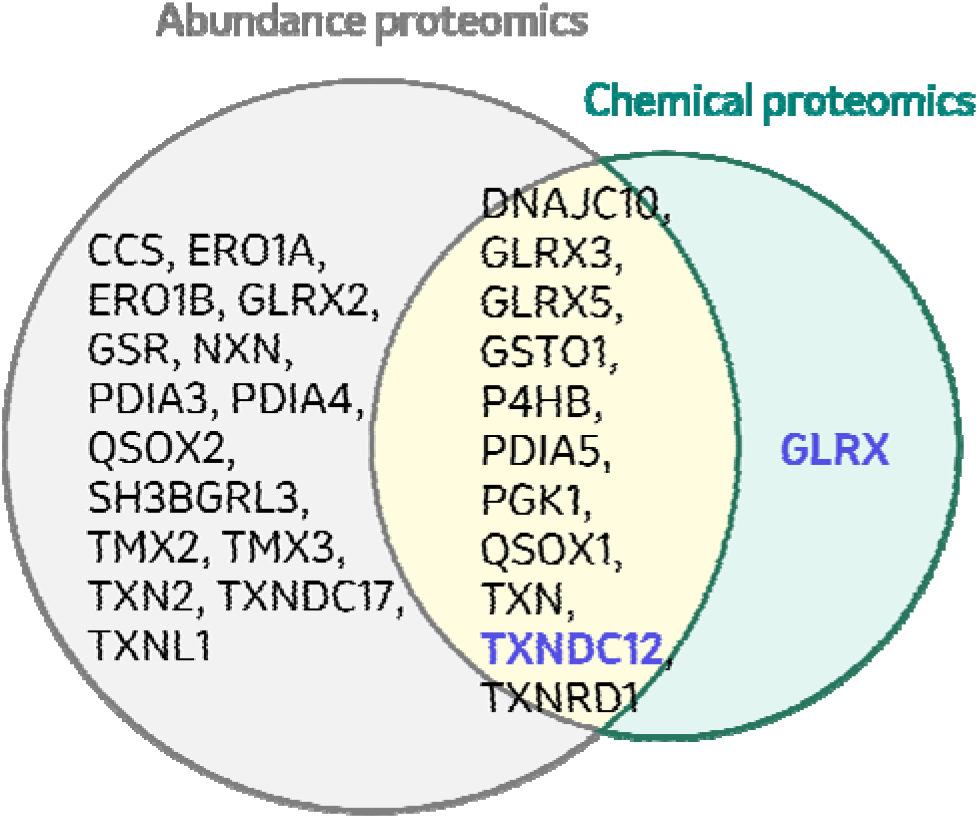
Overlap of disulfide oxidoreductase identifications between abundance-based proteomics and cysteine reactivity profiling (chemical proteomics). Active site cysteines of the enzymes highlighted in blue were found reactive towards the IA-DTB chemical reporter.

## CONCLUSIONS

In conclusion, we developed a chemical proteomics workflow to quantify cysteine reactivity in mAb-reducing biopharmaceutical samples. Previous studies on root causes of mAb disulfide reduction have relied on broad-spectrum inhibitors, recombinant enzymes and/or genetic knockdown.^9–11,14^ Unfortunately, it is difficult to pinpoint causative enzymes based on broad-spectrum inhibitors and genetic knockdown due to off-target and compensation effects. Biochemical reconstitution studies can demonstrate the activity of a recombinant enzyme towards a mAb substrate *in vitro*, but whether the same enzyme is indeed active *in situ* e.g., in HCCF often remains unclear. Alternatively, abundance proteomics enables unbiased profiling of HCP identity and abundance but not activity.^12,13^ In this regard, our cysteine reactivity profiling workflow probes the functional state of individual enzymes and other HCPs directly in biopharmaceutical process intermediates. It is also noteworthy that this workflow does not require external addition of NADPH, which is often done in biochemical reconstitution studies but would alter activity states of various redox enzymes. Furthermore, the workflow could be extended to profile cysteine oxidation and reactivity in cell culture process samples and critical organelles such as endoplasmic reticulum, ^35,36^ and even for biopharmaceutical stability samples where cysteine reactivity may play a critical role, such as cysteine-linked antibody-drug conjugates (ADC). Thus, the workflow should facilitate root-cause investigations and inspire novel control strategies for biopharmaceutical disulfide reduction.

## ASSOCIATED CONTENT

## Supporting Information

The Supporting Information is available free of charge.

Additional experimental details and results (PDF)

## Author Contributions

The manuscript was written through the contributions of all authors. All authors have given approval to the final version of the manuscript.

## Notes

The authors declare no competing financial interest.

## ACKNOWLEDGEMENT

T.T. thanks the MRL Postdoctoral Program. The authors thank Kaniz Fatema for proteomics sample preparation, and Hillary A. Schuessler for guidance and support.

## Notes

### Competing Interest Statement

The authors have declared no competing interest.

## REFERENCES

(1) Mullard, A. 2022 FDA Approvals. Nat. Rev. Drug Discov. 2023, 22 (2), 83–88. 10.1038/d41573-023-00001-3.

(2) Walsh, G.; Walsh, E. Biopharmaceutical Benchmarks 2022. Nat. Biotechnol. 2022, 40 (12), 1722–1760. 10.1038/s41587-022-01582-x.

(3) O’Flaherty, R.; Bergin, A.; Flampouri, E.; Mota, L. M.; Obaidi, I.; Quigley, A.; Xie, Y.; Butler, M. Mammalian Cell Culture for Production of Recombinant Proteins: A Review of the Critical Steps in Their Biomanufacturing. Biotechnol. Adv. 2020, 43 (May), 107552. 10.1016/j.biotechadv.2020.107552.

(4) Kunert, R.; Reinhart, D. Advances in Recombinant Antibody Manufacturing. Appl. Microbiol. Biotechnol. 2016, 100 (8), 3451–3461. 10.1007/s00253-016-7388-9.

(5) Liu, H.; Chumsae, C.; Gaza-Bulseco, G.; Hurkmans, K.; Radziejewski, C. H. Ranking the Susceptibility of Disulfide Bonds in Human IgG1 Antibodies by Reduction, Differential Alkylation, and LC-MS Analysis. Anal. Chem. 2010, 82 (12), 5219–5226. 10.1021/ac100575n.

(6) Feige, M. J.; Hendershot, L. M.; Buchner, J. How Antibodies Fold. Trends Biochem. Sci. 2010, 35 (4), 189–198. 10.1016/j.tibs.2009.11.005.

(7) Chung, W. K.; Russell, B.; Yang, Y.; Handlogten, M.; Hudak, S.; Cao, M.; Wang, J.; Robbins, D.; Ahuja, S.; Zhu, M. Effects of Antibody Disulfide Bond Reduction on Purification Process Performance and Final Drug Substance Stability. Biotechnol. Bioeng. 2017, 114 (6), 1264–1274. 10.1002/bit.26265.

(8) Ren, T.; Tan, Z.; Ehamparanathan, V.; Lewandowski, A.; Ghose, S.; Li, Z. J. Antibody Disulfide Bond Reduction and Recovery during Biopharmaceutical Process Development—A Review. Biotechnol. Bioeng. 2021, 118 (8), 2829–2844. 10.1002/bit.27790.

(9) Trexler-Schmidt, M.; Sargis, S.; Chiu, J.; Sze-Khoo, S.; Mun, M.; Kao, Y. H.; Laird, M. W. Identification and Prevention of Antibody Disulfide Bond Reduction during Cell Culture Manufacturing. Biotechnol. Bioeng. 2010, 106 (3), 452–461. 10.1002/bit.22699.

(10) Kao, Y. H.; Hewitt, D. P.; Trexler-Schmidt, M.; Laird, M. W. Mechanism of Antibody Reduction in Cell Culture Production Processes. Biotechnol. Bioeng. 2010, 107 (4), 622–632. 10.1002/bit.22848.

(11) Koterba, K. L.; Borgschulte, T.; Laird, M. W. Thioredoxin 1 Is Responsible for Antibody Disulfide Reduction in CHO Cell Culture. J. Biotechnol. 2012, 157 (1), 261–267. 10.1016/j.jbiotec.2011.11.009.

(12) Cura, A. J.; Xu, X.; Egan, S.; Aron, K.; Jenkins, L.; Hageman, T.; Huang, Y.; Chollangi, S.; Borys, M.; Ghose, S.; Li, Z. J. Metabolic Understanding of Disulfide Reduction during Monoclonal Antibody Production. Appl. Microbiol. Biotechnol. 2020, 104 (22), 9655–9669. 10.1007/s00253-020-10916-1.

(13) Park, S. Y.; Egan, S.; Cura, A. J.; Aron, K. L.; Xu, X.; Zheng, M.; Borys, M.; Ghose, S.; Li, Z.; Lee, K. Untargeted Proteomics Reveals Upregulation of Stress Response Pathways during CHO-Based Monoclonal Antibody Manufacturing Process Leading to Disulfide Bond Reduction. MAbs 2021, 13 (1). 10.1080/19420862.2021.1963094.

(14) Handlogten, M. W.; Zhu, M.; Ahuja, S. Glutathione and Thioredoxin Systems Contribute to Recombinant Monoclonal Antibody Interchain Disulfide Bond Reduction during Bioprocessing. Biotechnol. Bioeng. 2017, 114 (7), 1469–1477. 10.1002/bit.26278.

(15) O’Mara, B.; Gao, Z. H.; Kuruganti, M.; Mallett, R.; Nayar, G.; Smith, L.; Meyer, J. D.; Therriault, J.; Miller, C.; Cisney, J.; Fann, J. Impact of Depth Filtration on Disulfide Bond Reduction during Downstream Processing of Monoclonal Antibodies from CHO Cell Cultures. Biotechnol. Bioeng. 2019, 116 (7), 1669–1683. 10.1002/bit.26964.

(16) Batth, T. S.; Tollenaere, M. A. X.; Rüther, P.; Gonzalez-Franquesa, A.; Prabhakar, B. S.; Bekker-Jensen, S.; Deshmukh, A. S.; Olsen, J. V. Protein Aggregation Capture on Microparticles Enables Multipurpose Proteomics Sample Preparation. Mol. Cell. Proteomics 2019, 18 (5), 1027–1035. 10.1074/mcp.TIR118.001270.

(17) Bache, N.; Geyer, P. E.; Bekker-Jensen, D. B.; Hoerning, O.; Falkenby, L.; Treit, P. V; Doll, S.; Paron, I.; Müller, J. B.; Meier, F.; Olsen, J. V; Vorm, O.; Mann, M. A Novel LC System Embeds Analytes in Pre-Formed Gradients for Rapid, Ultra-Robust Proteomics. Mol. Cell. Proteomics 2018, 17 (11), 2284–2296. 10.1074/mcp.TIR118.000853.

(18) Skowronek, P.; Thielert, M.; Voytik, E.; Tanzer, M. C.; Hansen, F. M.; Willems, S.; Karayel, O.; Brunner, A. D.; Meier, F.; Mann, M. Rapid and In-Depth Coverage of the (Phospho-) Proteome With Deep Libraries and Optimal Window Design for Dia-PASEF. Mol. Cell. Proteomics 2022, 21 (9), 100279. 10.1016/j.mcpro.2022.100279.

(19) Tsukidate, T.; Stiving, A. Q.; Rivera, S.; Sahoo, A.; Madabhushi, S.; Li, X. DiaPASEF Enables High-Throughput Proteomic Analysis of Host Cell Proteins for Biopharmaceutical Process Development. Anal. Chem. 2024, 96 (32), 12999–13006. 10.1021/acs.analchem.4c00977.

(20) Cox, J.; Mann, M. MaxQuant Enables High Peptide Identification Rates, Individualized p.p.b.-Range Mass Accuracies and Proteome-Wide Protein Quantification. Nat. Biotechnol. 2008, 26 (12), 1367–1372. 10.1038/nbt.1511.

(21) Demichev, V.; Messner, C. B.; Vernardis, S. I.; Lilley, K. S.; Ralser, M. DIA-NN: Neural Networks and Interference Correction Enable Deep Proteome Coverage in High Throughput. Nat. Methods 2020, 17 (1), 41–44. 10.1038/s41592-019-0638-x.

(22) Kistner, F.; Grossmann, J. L.; Sinn, L. R.; Demichev, V. QuantUMS: Uncertainty Minimisation Enables Confident Quantification in Proteomics. bioRxiv 2023. 10.1101/2023.06.20.545604.

(23) Pham, T.V; Henneman, A.A.; Jimenez, C.R. IqL: An R Package to Estimate Relative Protein Abundances from Ion Quantification in DIA-MS-Based Proteomics. Bioinformatics 2020, 36 (8), 2611–2613. 10.1093/bioinformatics/btz961.

(24) Jeong, W.; Lee, D.; Park, S.; Rhee, S. G. BIOCHEMICAL PROPERTIES AND ROLE IN APOPTOSIS INDUCED BY ENDOPLASMIC. J. Biol. Chem. 2008, 283 (37), 25557–25566. 10.1074/jbc.M803804200.

(25) Hanschmann, E. M.; Godoy, J. R.; Berndt, C.; Hudemann, C.; Lillig, C. H. Thioredoxins, Glutaredoxins, and Peroxiredoxins-Molecular Mechanisms and Health Significance: From Cofactors to Antioxidants to Redox Signaling. Antioxidants Redox Signal. 2013, 19 (13), 1539– 1605. 10.1089/ars.2012.4599.

(26) Maurais, A. J.; Weerapana, E. ScienceDirect Reactive-Cysteine Profiling for Drug Discovery. Curr. Opin. Chem. Biol. 2019, 50 (1), 29–36. 10.1016/j.cbpa.2019.02.010.

(27) Weerapana, E.; Wang, C.; Simon, G. M.; Richter, F.; Khare, S.; Dillon, M. B. D.; Bachovchin, D. A.; Mowen, K.; Baker, D.; Cravatt, B. F. Quantitative Reactivity Profiling Predicts Functional Cysteines in Proteomes. Nature 2010, 468 (7325), 790–797. 10.1038/nature09472.

(28) Kuljanin, M.; Mitchell, D. C.; Schweppe, D. K.; Gikandi, A. S.; Nusinow, D. P.; Bulloch, N. J.; Vinogradova, E. V; Wilson, D. L.; Kool, E. T.; Mancias, J. D.; Cravatt, B. F.; Gygi, S. P. Reimagining High-Throughput Profiling of Reactive Cysteines for Cell-Based Screening of Large Electrophile Libraries. Nat. Biotechnol. 2021, 39 (5), 630–641. 10.1038/s41587-020-00778-3.

(29) Meier, F.; Brunner, A. D.; Frank, M.; Ha, A.; Bludau, I.; Voytik, E.; Kaspar-Schoenefeld, S.; Lubeck, M.; Raether, O.; Bache, N.; Aebersold, R.; Collins, B. C.; Röst, H. L.; Mann, M. DiaPASEF: Parallel Accumulation–Serial Fragmentation Combined with Data-Independent Acquisition. Nat. Methods 2020, 17 (12), 1229–1236. 10.1038/s41592-020-00998-0.

(30) Meier, F.; Brunner, A. D.; Koch, S.; Koch, H.; Lubeck, M.; Krause, M.; Goedecke, N.; Decker, J.; Kosinski, T.; Park, M. A.; Bache, N.; Hoerning, O.; Cox, J.; Räther, O.; Mann, M. Online Parallel Accumulation–Serial Fragmentation (PASEF) with a Novel Trapped Ion Mobility Mass Spectrometer. Mol. Cell. Proteomics 2018, 17 (12), 2534–2545. 10.1074/mcp.TIR118.000900.

(31) Yang, F.; Jia, G.; Guo, J.; Liu, Y.; Wang, C. Quantitative Chemoproteomic Profiling with Data-Independent Acquisition-Based Mass Spectrometry. J. Am. Chem. Soc. 2022, 144 (2), 901–911. 10.1021/jacs.1c11053.

(32) Corteselli, E. M.; Sharafi, M.; Hondal, R.; MacPherson, M.; White, S.; Lam, Y. W.; Gold, C.; Manuel, A. M.; van der Vliet, A.; Schneebeli, S. T.; Anathy, V.; Li, J.; Janssen-Heininger, Y. M. W. Structural and Functional Fine Mapping of Cysteines in Mammalian Glutaredoxin Reveal Their Differential Oxidation Susceptibility. Nat. Commun. 2023, 14 (1), 1–14. 10.1038/s41467-023-39664-2.

(33) Sevier, C. S. Erv2 and Quiescin Sulfhydryl OxidasesL: Erv-Domain Enzymes Associated with the Secretory Pathway. 2012, 16 (8), 800–808. 10.1089/ars.2011.4450.

(34) Komatsu, K.; Kumon, K.; Arita, M.; Onitsuka, M.; Omasa, T.; Yohda, M. Effect of the Disulfide Isomerase PDIa4 on the Antibody Production of Chinese Hamster Ovary Cells. J. Biosci. Bioeng. 2020, 130 (6), 637–643. 10.1016/j.jbiosc.2020.08.001.

(35) Sinharoy, P.; McFarland, K. S.; Majewska, N. I.; Betenbaugh, M. J.; Handlogten, M. W. Redox as a Bioprocess Parameter: Analytical Redox Quantification in Biological Therapeutic Production. Current Opinion in Biotechnology. Elsevier Ltd 2021, pp 49–54. 10.1016/j.copbio.2021.06.017.

(36) Bechtel, T. J.; Li, C.; Kisty, E. A.; Maurais, A. J.; Weerapana, E. Profiling Cysteine Reactivity and Oxidation in the Endoplasmic Reticulum. ACS Chem. Biol. 2020, 15 (2), 543–553. 10.1021/acschembio.9b01014.

