## Supporting Information for "Cysteine Reactivity Profiling Illuminates Monoclonal Antibody Disulfide Bond Reduction Mechanisms in Biopharmaceutical Process Intermediates"

**Table of Content**

Supplementary Figure 1	S2

Supplementary Table 1	S3

Supplementary Figure 2	S5

Supplementary Table 2	S6

Supplementary Script 1	S8

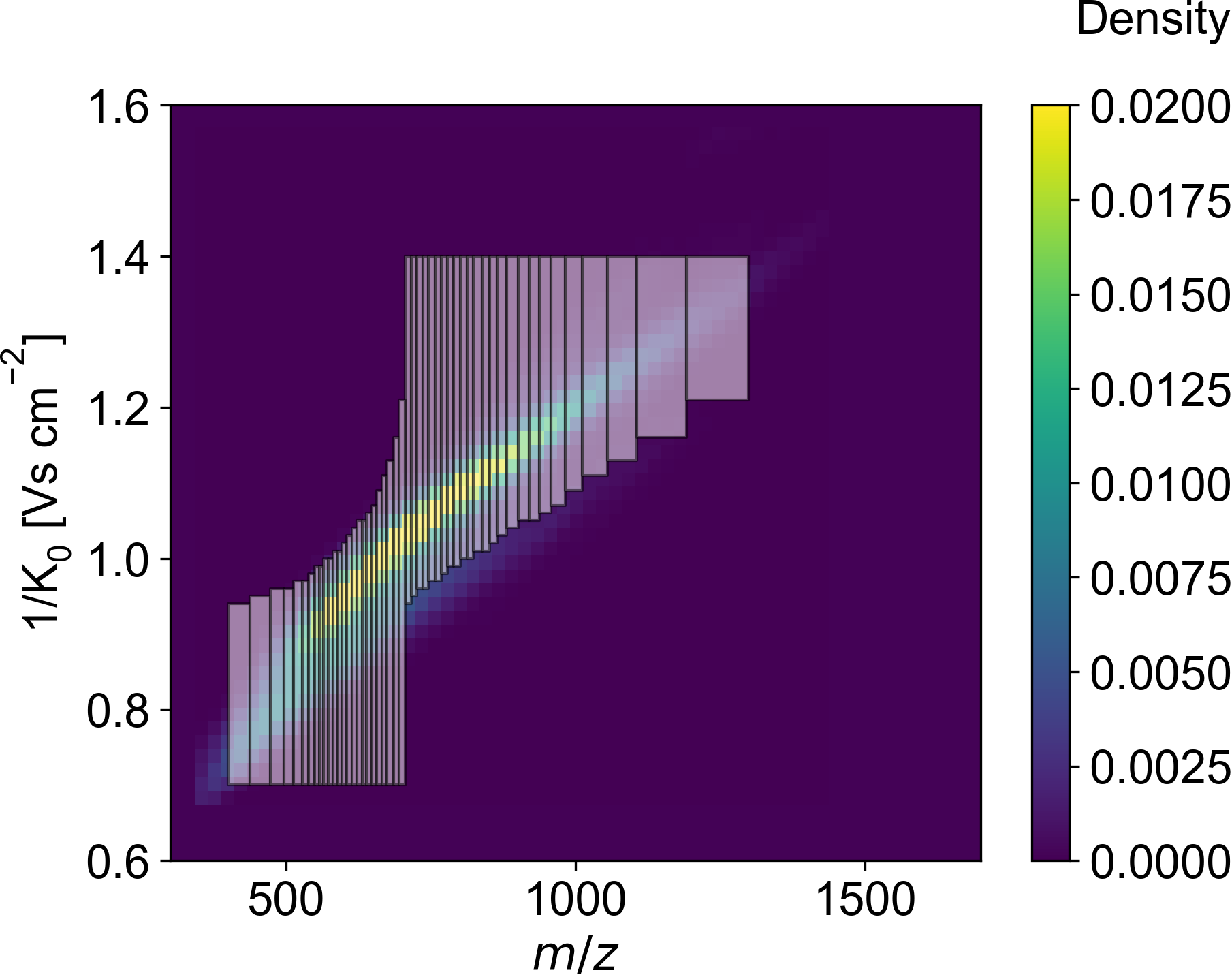

**Supplementary Figure 1.** diaPASEF method to target IA-DTB–modified precursors plotted as a kernel density distribution in the *m*/*z*–1/*K*_0_ space.

**Supplementary Table 1.** diaPASEF method to target IA-DTB–modified precursors. This table can be saved as a comma-separated text file and loaded in the Compass HyStar software to reproduce the method.

#----------------------------------------------------------------------------------------------------

\# MS Type | Cycle Id | 1/K0 Begin [Vs/cm2] | 1/K0 End [Vs/cm2] | Start Mass [m/z] | End Mass [m/z] | CE [eV]

#----------------------------------------------------------------------------------------------------

MS1, 0, - , - , - , - , -

PASEF, 1, 0.94, 1.4, 705.7, 716.38, -

PASEF, 1, 0.7, 0.94, 400.19, 437.89, -

PASEF, 2, 0.95, 1.4, 716.38, 725.35, -

PASEF, 2, 0.7, 0.95, 437.89, 473.25, -

PASEF, 3, 0.96, 1.4, 725.35, 736.87, -

PASEF, 3, 0.7, 0.96, 473.25, 495.94, -

PASEF, 4, 0.96, 1.4, 736.87, 746.41, -

PASEF, 4, 0.7, 0.96, 495.94, 512.57, -

PASEF, 5, 0.97, 1.4, 746.41, 757.39, -

PASEF, 5, 0.7, 0.97, 512.57, 527.95, -

PASEF, 6, 0.97, 1.4, 757.39, 767.69, -

PASEF, 6, 0.7, 0.97, 527.95, 538.3, -

PASEF, 7, 0.98, 1.4, 767.69, 778.88, -

PASEF, 7, 0.7, 0.98, 538.3, 548.97, -

PASEF, 8, 0.99, 1.4, 778.88, 789.37, -

PASEF, 8, 0.7, 0.99, 548.97, 557.81, -

PASEF, 9, 0.99, 1.4, 789.37, 801.05, -

PASEF, 9, 0.7, 0.99, 557.81, 565.31, -

PASEF, 10, 1.0, 1.4, 801.05, 812.42, -

PASEF, 10, 0.7, 1.0, 565.31, 572.81, -

PASEF, 11, 1.0, 1.4, 812.42, 823.89, -

PASEF, 11, 0.7, 1.0, 572.81, 580.81, -

PASEF, 12, 1.01, 1.4, 823.89, 838.44, -

PASEF, 12, 0.7, 1.01, 580.81, 588.96, -

PASEF, 13, 1.01, 1.4, 838.44, 851.5, -

PASEF, 13, 0.7, 1.01, 588.96, 597.06, -

PASEF, 14, 1.02, 1.4, 851.5, 864.92, -

PASEF, 14, 0.7, 1.02, 597.06, 604.81, -

PASEF, 15, 1.03, 1.4, 864.92, 881.44, -

PASEF, 15, 0.7, 1.03, 604.81, 614.01, -

PASEF, 16, 1.04, 1.4, 881.44, 901.44, -

PASEF, 16, 0.7, 1.04, 614.01, 622.62, -

PASEF, 17, 1.05, 1.4, 901.44, 919.46, -

PASEF, 17, 0.7, 1.05, 622.62, 631.37, -

PASEF, 18, 1.05, 1.4, 919.46, 938.1, -

PASEF, 18, 0.7, 1.05, 631.37, 638.85, -

PASEF, 19, 1.06, 1.4, 938.1, 957.0, -

PASEF, 19, 0.7, 1.06, 638.85, 648.65, -

PASEF, 20, 1.07, 1.4, 957.0, 982.46, -

PASEF, 20, 0.7, 1.07, 648.65, 656.86, -

PASEF, 21, 1.09, 1.4, 982.46, 1012.03, -

PASEF, 21, 0.7, 1.09, 656.86, 665.84, -

PASEF, 22, 1.11, 1.4, 1012.03, 1055.5, -

PASEF, 22, 0.7, 1.11, 665.84, 674.33, -

PASEF, 23, 1.13, 1.4, 1055.5, 1105.51, -

PASEF, 23, 0.7, 1.13, 674.33, 685.35, -

PASEF, 24, 1.16, 1.4, 1105.51, 1191.05, -

PASEF, 24, 0.7, 1.16, 685.35, 694.85, -

PASEF, 25, 1.21, 1.4, 1191.05, 1298.6, -

PASEF, 25, 0.7, 1.21, 694.85, 705.7, -

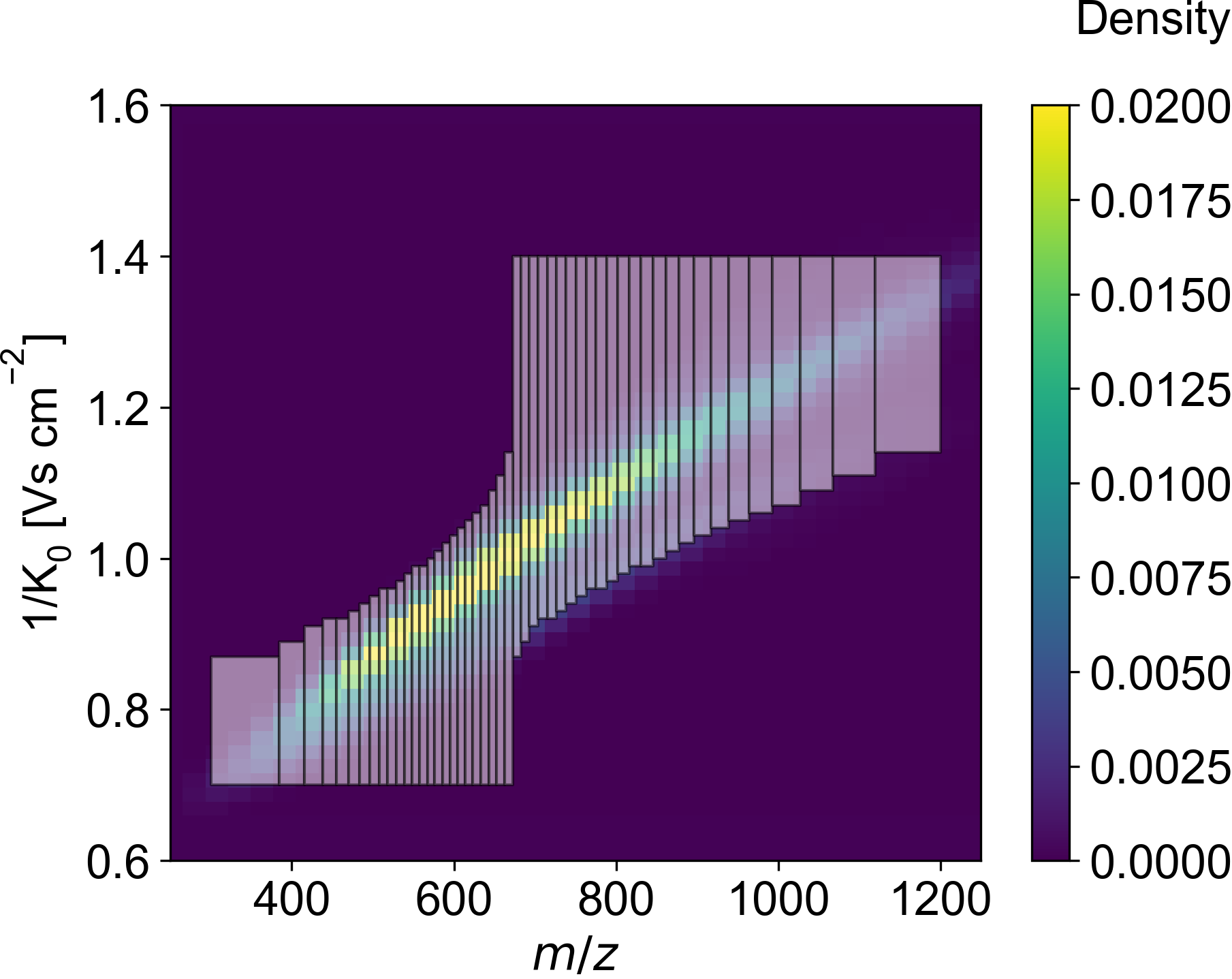

**Supplementary Figure 2.** diaPASEF method to target HeLa protein digest–derived precursors plotted as a kernel density distribution in the *m*/*z*–1/*K*_0_ space.

**Supplementary Table 2.** diaPASEF method to target HeLa protein digest–derived precursors. This table can be saved as a comma-separated text file and loaded in the Compass HyStar software to reproduce the method.

#----------------------------------------------------------------------------------------------------

\# MS Type | Cycle Id | 1/K0 Begin [Vs/cm2] | 1/K0 End [Vs/cm2] | Start Mass [m/z] | End Mass [m/z] | CE [eV]

#----------------------------------------------------------------------------------------------------

MS1, 0, - , - , - , - , -

PASEF, 1, 0.87, 1.4, 672.35, 682.67, -

PASEF, 1, 0.7, 0.87, 300.5, 384.46, -

PASEF, 2, 0.89, 1.4, 682.67, 692.89, -

PASEF, 2, 0.7, 0.89, 384.46, 415.71, -

PASEF, 3, 0.91, 1.4, 692.89, 703.68, -

PASEF, 3, 0.7, 0.91, 415.71, 437.84, -

PASEF, 4, 0.92, 1.4, 703.68, 715.35, -

PASEF, 4, 0.7, 0.92, 437.84, 454.75, -

PASEF, 5, 0.92, 1.4, 715.35, 726.33, -

PASEF, 5, 0.7, 0.92, 454.75, 470.24, -

PASEF, 6, 0.93, 1.4, 726.33, 738.37, -

PASEF, 6, 0.7, 0.93, 470.24, 483.55, -

PASEF, 7, 0.94, 1.4, 738.37, 750.85, -

PASEF, 7, 0.7, 0.94, 483.55, 496.24, -

PASEF, 8, 0.95, 1.4, 750.85, 762.99, -

PASEF, 8, 0.7, 0.95, 496.24, 507.59, -

PASEF, 9, 0.96, 1.4, 762.99, 775.4, -

PASEF, 9, 0.7, 0.96, 507.59, 518.3, -

PASEF, 10, 0.96, 1.4, 775.4, 788.38, -

PASEF, 10, 0.7, 0.96, 518.3, 529.26, -

PASEF, 11, 0.97, 1.4, 788.38, 802.39, -

PASEF, 11, 0.7, 0.97, 529.26, 538.74, -

PASEF, 12, 0.98, 1.4, 802.39, 816.41, -

PASEF, 12, 0.7, 0.98, 538.74, 548.58, -

PASEF, 13, 0.99, 1.4, 816.41, 830.45, -

PASEF, 13, 0.7, 0.99, 548.58, 557.83, -

PASEF, 14, 0.99, 1.4, 830.45, 845.88, -

PASEF, 14, 0.7, 0.99, 557.83, 567.3, -

PASEF, 15, 1.0, 1.4, 845.88, 861.39, -

PASEF, 15, 0.7, 1.0, 567.3, 576.36, -

PASEF, 16, 1.01, 1.4, 861.39, 877.9, -

PASEF, 16, 0.7, 1.01, 576.36, 585.82, -

PASEF, 17, 1.02, 1.4, 877.9, 896.41, -

PASEF, 17, 0.7, 1.02, 585.82, 595.29, -

PASEF, 18, 1.03, 1.4, 896.41, 917.44, -

PASEF, 18, 0.7, 1.03, 595.29, 604.32, -

PASEF, 19, 1.04, 1.4, 917.44, 938.96, -

PASEF, 19, 0.7, 1.04, 604.32, 613.82, -

PASEF, 20, 1.05, 1.4, 938.96, 963.91, -

PASEF, 20, 0.7, 1.05, 613.82, 623.29, -

PASEF, 21, 1.06, 1.4, 963.91, 992.86, -

PASEF, 21, 0.7, 1.06, 623.29, 632.81, -

PASEF, 22, 1.07, 1.4, 992.86, 1027.07, -

PASEF, 22, 0.7, 1.07, 632.81, 642.81, -

PASEF, 23, 1.09, 1.4, 1027.07, 1067.58, -

PASEF, 23, 0.7, 1.09, 642.81, 652.31, -

PASEF, 24, 1.11, 1.4, 1067.58, 1119.14, -

PASEF, 24, 0.7, 1.11, 652.31, 662.35, -

PASEF, 25, 1.14, 1.4, 1119.14, 1199.89, -

PASEF, 25, 0.7, 1.14, 662.35, 672.35, -

**Supplementary Script 1.** R script to format MaxQuant evidence and msms files to feed to DIA-NN.

```
library(readr)
library(dplyr)
library(stringr)
library(tidyr)
library(tibble)

# (Protein names, Gene names, Proteotypicity) --> *,*,*
# Decoy: filter out Reverse and convert NA to 0
# Extract Fragment charge, ion type and series number from Matches
# Extract Fragment mass and intensity from Masses and Intensities
# Neutral loss level none --> noloss, H2O, NH3
# (Q-value, Elution group identifier, Exclude fragment indicator) --> *,*,*
# Ion mobility from the evidence file

# Extract calibrated retention time and ion mobility from the evidence file
evi_ <- evi %>%
  filter(is.na(Reverse), !is.na(`Modified sequence`)) %>%
  select(
    `Raw file`,
    `Modified sequence`,
    `Charge`,
    `Retention time`,
    `Calibrated retention time`,
    `Calibrated 1/K0`
    )

# Select relevant columns from the msms file
msms_ <- msms %>%
  filter(is.na(Reverse)) %>%
  select(
    `Raw file`,
    `Modified sequence`,
    `Charge`,
    `m/z`,
    `Retention time`,
    `Masses`,
    `Intensities`,
    `Proteins`,
    `Reverse`,
    `Matches`,
    `Neutral loss level`
    )

out <- msms_ %>%
  left_join(evi_) %>%
  mutate(
    Reverse = 0,
    Fragment = str_split(Matches, ";"),
    Mass = str_split(Masses, ";"),
    Intensity = str_split(Intensities, ";")
    ) %>%
  unnest(c(Fragment, Mass, Intensity)) %>%
  mutate(
    `Fragment ion type` = str_extract(Fragment, "[abcxyz]"),
    `Fragment ion type` = if_else(`Fragment ion type` %in% c("a", "b", "c"), "b", "y"),
    `Fragment series number` = str_extract(Fragment, "[:digit:]+"),
    `Fragment ion charge` = str_extract(Fragment, "[:digit:](?=\\+)"),
    `Fragment ion charge` = if_else(!is.na(`Fragment ion charge`), `Fragment ion charge`, "1"),
    `Neutral loss level` = str_extract(Fragment, "H2O|NH3"),
    `Neutral loss level` = if_else(!is.na(`Neutral loss level`), `Neutral loss level`, "noloss")
    ) %>%
  select(-c(Matches, Fragment, Masses, Intensities))

write_tsv(out, "evimsms.tsv")

#In DIA-NN, add the following command to Additional options:
# --library-headers
# Modified sequence,Charge,m/z,Calibrated retention time,Mass,Intensity,Proteins,*,*,*,Reverse,Fragment ion charge,Fragment ion type,Fragment series number,Neutral loss level,*,*,*,Calibrated 1/K0
```
